# DEAD-box RNA helicase DDX3X maintains the homeostasis of the Zika virus translation-replication cycle

**DOI:** 10.64898/2026.03.27.714787

**Authors:** Tomás Hernández-Díaz, Alonso Gonzalez-Poblete, Sebastián Giraldo-Ocampo, Aaron Oyarzún-Arrau, Cecilia Rojas-Fuentes, Lucía Cortesi-Patiño, Fernando Carrasco-Gálvez, Delia López-Palma, Mónica L. Acevedo, Barbara Rojas-Araya, Marcelo López-Lastra, Aracelly Gaete-Argel, Fernando Valiente-Echeverria, Tommaso Cupido, Matías Zúñiga-Bustos, Ricardo Soto-Rifo

## Abstract

DDX3X is a cellular DEAD-box ATP-dependent RNA helicase known to play pivotal roles during the replication cycle of different viruses including some flaviviruses. Whether DDX3X plays a role during replication of Zika virus (ZIKV), a mosquito-borne flavivirus with a broad tissue tropism, has not been explored in detail. Here, we show that DDX3X is required for efficient ZIKV replication in a human microglia cell line but no other cell lines. Mechanistically, we provide evidence showing that DDX3X is recruited to the viral replication compartments where it binds to the 5’UTR of the ZIKV RNA and promotes viral protein synthesis in an ATP-dependent manner. We also show that DDX3X binds to the viral RNA-dependent RNA polymerase NS5 and interferes with viral RNA synthesis in an ATP-independent manner. Such an effect was not observed during replication of Dengue 2 virus in human microglia, revealing specific roles for DDX3X in the regulation of the translation-replication cycle of ZIKV in human microglia.

**IMPORTANCE:** Zika virus emerged as a major threat to humans due to its pandemic potential and its association with birth defects and neurological complications. The lack of available vaccines or specific treatments makes the understanding of Zika virus-host interactions a priority. Here, we demonstrate a mechanism by which DDX3X, a cellular ATP-dependent RNA helicase, promotes Zika virus replication by regulating the translation-replication cycle in a human microglia cell line. Considering the threat of this virus to the human population and the wide range of RNA viruses that usurp this host protein to accomplish the replication cycle, DDX3X continuously rises as a potential and valuable target for pharmacological intervention.

## INTRODUCTION

Zika virus (ZIKV) is a mosquito-borne flavivirus transmitted either through the bite of *Aedes* mosquitoes or directly between humans through sexual contact, blood transfusions or from mother to child during pregnancy (1–3). In January 2016, the World Health Organization (WHO) declared ZIKV as a global threat to public health given its ability to cause neurological disorders such as microcephaly in newborns and Guillain-Barré syndrome in adults (4, 5). Although preferentially neurotropic, ZIKV has a broad tropism being able to persist in different tissues and fluids (2). The viral RNA (vRNA) genome consists of an 11-kb single-stranded positive-sense molecule encoding a polyprotein surrounded by untranslated regions (UTR) at each end (6–8). The 5’UTR contains a cap0 and cap1 structure and the 3’UTR lacks a poly-A tail. The untranslated regions at both ends of the viral genome adopt highly conserved structures important in the regulation of translation and replication of the vRNA (9, 10). ZIKV vRNA translation takes place in the endoplasmic reticulum (ER) at the viral replication compartments (VRC) leading to the synthesis of a polyprotein that is processed into individual structural and non-structural proteins by the viral protease/helicase NS3 and cellular proteases of the ER (7, 11). The vRNA is also used as the template for the synthesis of a negative strand by the RNA-dependent RNA polymerase (RdRp) NS5, which then is used for the generation of new copies of vRNA that will be packaged into newly produced virions. As such, NS5 has been shown to inhibit *de novo* vRNA translation initiation to promote RNA synthesis (12, 13). However, whether host factors are involved in the regulation of these processes is not well understood.

DDX3X, a member of the DEAD (Asp-Glu-Ala-Asp)-box helicases family, is ubiquitously expressed in different tissues and displays ATP-dependent RNA helicase activity (14, 15). As such, this cellular RNA helicase plays important roles in various biological processes, including translation initiation of selected mRNA, stress granules assembly, innate immune response signaling, and NLRP3 inflammasome activation (15–19). Due to its multifaceted functions, different viruses including important human pathogens such as human immunodeficiency virus type 1 (HIV-1), hepatitis C virus (HCV), SARS-CoV-2, Japanese Encephalitis virus (JEV) and West Nile virus (WNV) usurp DDX3X to promote their replication cycles (20–29).

In this work, we characterized the role of DDX3X in ZIKV replication during infection in a human microglia cell line, an important cell target associated to viral pathogenesis in the CNS (30–35). Our data uncover a critical role of DDX3X in regulating the homeostasis of the vRNA translation-replication cycle by preserving the synthesis of viral proteins required for efficient production of the viral progeny. As such, we also show that DDX3X is recruited to the VRC in the endoplasmic reticulum, binds the viral RNA at the 5’UTR and ensures the synthesis of viral proteins in an ATP-dependent manner. DDX3X also interacts with the RdRp NS5 and interferes with the synthesis of both strands during vRNA replication. Together, our results provide mechanistic insights on how ZIKV recruits the host RNA helicase DDX3X to maintain the homeostasis of the translation-replication cycle.

## MATERIAL & METHODS

### Cell culture and virus infection

Human microglia C20, HEK293T and VeroE6 cell lines were maintained in DMEM culture media (Cat. SH30081.02, Cytiva lifescience) supplemented with 10% FBS (Cat. F0926, Merck), antibiotic-antimycotic 1% (Cat. A5955, Merck), L-glutamine 1% (Cat. A29168-01, Gibco). A549 cells were maintained in DMEM/F12 culture media (Cat. SH30023.02, Cytiva lifescience) supplemented with 10% FBS, antibiotic-antimycotic 1%, L-glutamine 1%. All mammalian cells were cultured at 37°C and 5% CO_2_. *Aedes albopictus* mosquito cell line C6/36 (ATCC-CRL1660, kindly donated by Paquita García, National Health Institute, Perú), were maintained in Leibovitz L-15 medium (Cat. SH30525.02, Cytiva lifescience), supplemented with 10% FBS, antibiotic-antimycotic 1%, L-glutamine 1% and were cultured at 28°C without CO_2_.

For Zika virus and Dengue virus infection, 3×10^5^ human microglia C20 cells were plated in 60 mm plates and grown until the next day in 4 mL of DMEM 10% FBS culture media. The next day, viral infection was carried out by adding the viral stock (MOI 3) to cells in DMEM 2% FBS for 1 hour for viral adsorption. Then, cells were washed with sterile 1X PBS (Cat. 46-013-CM, Corning) and 2.5 mL of DMEM 10% FBS culture media was added.

### Viral production, propagation, and titration

Viral stocks were obtained from a reverse genetics system for ZIKV described by Mutso et al (2017)(^36^). Briefly, the plasmid pCC1-SP6-ZIKV (kindly provided by Dr. Andres Merits, University of Tartus) that encodes a complete viral genome (ZIKV isolate BeH819015, GenBank accession n° KX197192) was used as a template for in vitro transcription using the mMessage mMachine^TM^ SP6 transcription kit (Cat. AM1340, Invitrogen) following the recommendations of the supplier to produce an infective RNA (iRNA)(36). 5 µg of iRNA were used to electroporate 3×10^6^ VeroE6 cells using the Neon^TM^ transfection system (ThermoFisher Scientific). The virus was recovered three days after electroporation from the cell supernatant (passage 0) and was mixed with one volume of FBS. Then, the virus stock was stored at -80°C until use. Viral stocks were used to infect VeroE6 cells successively and obtain up to 5 more virus passages. For dengue virus (DENV-2), we used the serotype dengue virus 2 strain NGC (kindly donated by Marcelo López-Lastra, ATCC VR-1584). This virus was propagated in *Aedes albopictus* C6/36 cells at 28 °C without CO_2_ (37). Viral titers of each stock were determined by a plate lysis assay. Briefly, VeroE6 monolayers were infected with serial dilutions of viral stocks. After 1 hour of viral adsorption, the culture media was replaced by a semi-viscous medium (DMEM 10% FBS and carboxymethylcellulose 0.75%). After 5 to 6 days post-infection, the cells were fixed with 37% formalin overnight at room temperature, then the formalin and medium were removed from the wells, and cells were stained with 1% crystal violet (Sigma-Aldrich) for 2 hours. Finally, the crystal violet (Cat. 1.04137, Sigma-Aldrich) was washed, and the plaques formed in each well were manually counted (36–38).

### DDX3X targeting

For DDX3X silencing, C20 microglia were seeded at 70% confluency and transfected with 10 nM of a control siRNA (sc-37007, Santa Cruz Biotechnologies) or with a previously described siRNA targeting the DDX3X transcript (5’–GAUGCUGGCUCGUGAUUUCUdTdT–3’)(23, 39). For this, Lipofectamine^TM^ RNAiMAX Transfection Reagent (Cat. 13778150, Invitrogen) was used following the recommendations of the manufacturer and 24 hours after transfection the cells were infected with ZIKV for 24 hours as described above. Silencing efficiency was evaluated by Western blot using anti-DDX3X antibody (Cat. 11115-1-AP, Proteintech).

Pharmacological inhibition of DDX3X was carried out with the specific ATP-binding inhibitor RK-33 (Cat. 10-4839, Focus Biomolecules)(40). For this purpose, a stock solution of RK-33 was prepared in 100% DMSO and diluted in DMEM 10% FBS media according to the working concentrations. C20 microglia cells were seeded at 70% confluency and after 1 hour of viral adsorption, the medium was replaced with fresh medium containing the different concentrations of RK-33 and medium with 2% DMSO was used as a control (40).

### RNA extraction and RT-qPCR

Cellular RNA extraction and RT-qPCR were performed as we have previously described (41). Briefly, RNA was extracted from cell lysates using TRIzol® Reagent (Cat. 15596026, Invitrogen) as indicated by the manufacturer and total RNA was treated with Turbo DNase (Cat. AM2238, Invitrogen). Then, we used 300 ng of RNA to reverse-transcription reaction using the High-Capacity cDNA Reverse Transcription Kit (Cat. 4368814, Invitrogen) following the supplier’s indications. The cDNA was diluted 10 times in nuclease-free water, and qPCR was performed in duplicate using the Brilliant II SYBR® Green QPCR Master Mix (Cat. 600828, Agilent Technologies). The amplification reaction was carried out using the AriaMx Real-Time PCR System (Agilent Technologies). The relative quantity of transcript levels was normalized to 18S rRNA using the ΔΔCt method previously described (42) using the following primers: 18S rRNA Fw 5’*CCCTGCCCTTTGTACACACC’*3 – Rv 5’*CGATCCGAGGGCCTCACTA*’3, DDX3X Fw 5’*GAAGCTACTAAAGGTTTCTAC*’3 – Rv 5’*TCTCAACATCACTGAAACTTTC*’3, IFN-β Fw 5’*TCTAGCACTGGCTGGAATGAGACT*’3 – Rv 5’*TGGCCTTCAGGTAATGCAGAATCC*’3, ZIKV vRNA Fw 5*’GTCTTGGAACATGGAGG’3* – Rv 5*’TTCACCTTGTGTTGGGC’*3, ZIKV (-)RNA Fw 5*’ CCTCCATGTTCCAAGAC’3* – Rv 5’*GCCCAACACAAGGTGAA’3,* DENV-2 vRNA Fw 5*’GCTGCTGACAAAACCTTGGG’3* – Rv 5*’ TCGGTTCTTGGGTTCTCGTG’*3, pCC1-SP6 ZIKV 5’UTR Fw 5*’AACGACGGCCAGTGAATTC*’3 – Rv 5*’ACCGGTAGACGGGCTACTCCGCG’*3, pCC1-SP6 ZIKV-NanoLuciferase Fw 5’*AGTGAATTCATTTAGGTGACA*’3 - Rv 5’AGACCCATGGATTCATCACGCCAGAATGCGTTCGCACAGCCGCCAGCC’3.

### Western blot

Proteins were extracted from cell cultures using RIPA lysis buffer (150 mM NaCl, 50 mM Tris HCl, 1% NP40, 0.1% SDS, 0.5% sodium deoxycolate, and protease inhibitor cocktail, Roche) and quantified using the Pierce™ BCA Protein Assay Kit (Cat. 23227, ThermoFisher Scientific). For SDS-PAGE electrophoresis, 40 µg of total protein was loaded in 5%-15% gradient gels. Gel electrophoresis was carried out at 120 V for 2 hours and the gels were transferred on a nitrocellulose membrane (Cat. 10600002, Cytiva Lifesciences) at a constant 400 mA for 2 hours. The membranes were blocked with 5% Blotting-Grade Blocker (Cat. 1706404, Bio-Rad) diluted in TBS-T 1X, and after 3 washes with TBS-T 1X, membranes were incubated overnight with the corresponding primary antibodies at 4°C. The antibodies used were anti-β-actin (sc-477778, mouse, 1:5000, Santa Cruz Biotechnologies), anti-DDX3X (sc-365768, mouse, 1:3000, Abcam), anti-RIG-I (20566-1-AP, rabbit, 1:1000, Proteintech), DENV-2 anti-NS3 (Cat. 124252, rabbit, 1:3000, Genetex), ZIKV anti-NS3 and ZIKV anti-NS5 (rabbit, 1:3000, gently donated by Dr. Andres Merits, University of Tartu, Estonia). After incubation, the membranes were washed 3 times with TBS-T 1X and then incubated for 2 hours at room temperature with HRP-conjugated secondary antibody (Cat. 111-035-003, 1:5000, Jackson Immunoresearch). Finally, for the detection and digitalization of images, the membrane was incubated with the chemiluminescent solution for Western blot EZ-ECL (ThermoFisher Scientific) and registered with a Q9 Alliance UVITEC-Cambridge digital photodocumenter. Densitometric analyses were performed using the Fiji-Image-J software (43).

### Fluorescence *in situ* hybridization, immunofluorescence and confocal microscopy

For viral genome detection, a specific digoxigenin-labeled RNA probe was synthesized. Briefly, a PCR with primers containing the T3 RNA polymerase promoter sequence was performed using the pCC1-SP6 ZIKV plasmid as a template. An *in vitro* transcription was conducted using the PCR product in transcription reaction buffer 1X with T3 RNA polymerase (Cat. P2083, Promega Corporation), 5 mM DTT, digoxigenin RNA labeling mix 1X (Cat. 11277073910, Roche) and 20 Units of RNAsin (Cat. N2511, Promega Corporation). The mixture was incubated for 2 hours at 37°C. RNA was precipitated in the presence of 2.5 M LiCl and 2 volumes of ethanol at -80°C for 1 hour. Then, *in vitro* transcribed RNA was centrifuged at 16000g for 30 min at 4°C, washed with 70% ethanol, and resuspended in 50 µL of nuclease-free water. RNA integrity was monitored by electrophoresis on agarose gels. Finally, RNA was treated with Turbo DNase and fragmented with RNA Fragmentation Reagents following manufacturer’s indications (Cat. AM8740, Ambion).

To perform RNA fluorescent *in situ* hybridization coupled to indirect immunofluorescence (RNA FISH-IFI), C20 microglia cells were seeded on coverslips. After infection and treatments, the cells were fixed with 4% paraformaldehyde for 20 minutes, incubated with glycine for 10 minutes and permeabilized with 0.2% Triton X-100 in 1X PBS for 5 minutes. Fifty nanograms of digoxigenin-labeled ZIKV RNA probe were hybridized overnight at 37°C in a solution containing 2 mM VRC, 0.02% BSA, 50% formamide and 1.5 µg/µL tRNA. The coverslips were then washed with a 50% formamide-0.2X SSC solution for 30 minutes at 50°C and then incubated with primary antibodies in antibody solution (2X Saline-Sodium Citrate, 8% formamide, 2 mM Vanadyl Ribonucleoside Complex, 0.02% BSA) for 1 hour at 37°C. Primary antibodies used were: anti-digoxigenin (sheep 1:200, Cat. 11333089001, Roche) and anti-NS3, anti-NS5 (rabbit 1:100, gently donated by Dr. Andres Merits). After incubation, unbound primary antibodies were washed 3 times with 1X PBS, then incubated with the respective secondary antibodies (1:500) or Alexa Fluor 647 Phalloidin for IFI (A22287, ThermoFisher Scientific). Finally, the coverslips were stained with DAPI (Cat D1306, Invitrogen), washed twice with PBS 1X then once with nuclease-free water and mounted on slides using aqueous mounting solution (Cat. F4680, Merck). Images (2D) were obtained with an inverted OLYMPUS IX73 epifluorescence microscope (40X objective) and 2D images and Z-stacks were obtained with a Zeiss LSM 800 Confocal Microscope (Zeiss) and processed using FIJI/ImageJ and Imaris software. For Mean Fluorescence Intensity (MFI) analysis, laser intensity was adjusted to avoid pixel saturation and threshold signal was manually set for each channel based on background signal/control conditions. Colocalization analysis expressed as the fraction of overlap (Mander’s overlap coefficients) of two selected channels was performed using BIOP JACoP plugin with an automated threshold defined by Costes method.

### DDX3X cross-linking and immunoprecipitation RT-qPCR (DDX3X-CLIP RT-qPCR)

C20 microglia were infected with ZIKV for 24 hours as described above. After infection, the cells were fixed with 0.1% formaldehyde for 10 minutes followed by quenching with quenching buffer (2M glycine, 10 mM Tris HCl pH 7.5) and then lysed using lysis buffer (150 mM NaCl, 50 mM Tris HCl, 1% NP40, 0.1% Triton X-100, 10 mM EDTA and protease inhibitor cocktail, Roche). For C20 microglia infected and treated with DMSO or 3 µM of RK-33, the cells were washed with PBS 1X once, crosslinked with 150 mJ/cm2 of UV-light at 254nm, and then trypsinized for 5-10 min at 37°C. Cells were collected, pelleted, washed once with PBS 1X, and then lysed using RIPA buffer (150 mM NaCl, 50 mM Tris HCl, 1% NP40, 0.1% SDS, 0.5% sodium deoxycolate, and protease inhibitor cocktail, Roche). Cell lysates were quantified using the Pierce BCA Protein Assay. Then, 5% of the lysate was set aside as the input fraction. The cell lysate was split into two fractions (300 ug each) and incubated overnight under constant shaking at 4°C with 0.5 µg of rabbit anti-DDX3X antibody (mouse, Santa Cruz Biotechnologies and rabbit, Proteintech) or 0.5 µg of normal rabbit IgG antibody (Cat. 12-370, Millipore) as a control. The DDX3X-antibody complexes were recovered using Dynabeads Protein G (Cat. 10003D, Invitrogen) previously washed twice with RIPA buffer. Then, DDX3X-antibody-beads complexes were washed twice with RIPA buffer, twice with RIPA-S buffer (500 mM NaCl, 50 mM Tris HCl, 1mM EDTA, 1% NP40, 0.1% SDS, 0.5% sodium deoxycolate, and protease inhibitor cocktail, Roche), and twice with Low wash buffer (150 mM NaCl, 25 mM Tris HCl, 5mM EDTA, 0.5% NP40, and protease inhibitor cocktail, Roche). 60% Input and bead samples were treated with 1.6 Units of proteinase K (NEB, P8107S) at 56°C for 1 hour. Input and 60% of bead samples were treated with 1.6 Units of proteinase K (P8107S, NEB) at 56°C for 1 hour. Then, total RNA was isolated from each fraction using TRIzol. 50 ng of RNA was used to perform RT-qPCR using KAPA SYBR® FAST One-Step (Cat. SFUKB, Roche) following the manufacturer’s instructions. In parallel, the remaining 40% of the samples were prepared for Western blot to check the immunoprecipitation of of DDX3X(39, 44).

### DDX3X (132–607) purification

Recombinant DDX3X was expressed using a plasmid containing amino acids 132-607 fused to a His6 tag and a maltose binding protein (45). The recombinant protein was expressed and purified from 2 liters of E. coli BL21 induced with Isopropyl-1-thio-β-d-galactopyranose (IPTG) for 16 hours at 16°C while shaking. Subsequently, the cells were harvested by centrifugation at 1000g for 10 minutes at 4°C and lysed with lysis buffer (500 mM NaCl, 50 mM Tris HCl, 5 mM MgCl_2_, 10% glycerol, 1 mM imidazole, 3 mM Dithiothreitol, 0.4 mM Phenylmethylsulfonyl fluoride (PMSF), 1X protease inhibitor, Roche). Cell lysate was clarified by ultracentrifugation using a Sorvall T45 rotor T647.5 at 30000g for 45 minutes at 4°C. The clarified lysate was passed through a nickel affinity column using the lysis buffer without PMSF and with imidazole at 500 mM as an elution buffer. Then, the sample was purified using ion exchange chromatography with an elution buffer (1 M NaCl, 20 mM HEPES pH 7.4, 5 mM MgCl_2_, 10% glycerol, 2 mM DTT). Enriched fractions from Heparin affinity chromatography were concentrated and injected into a Superdex 75 Increase 10/300GL column (Cytiva) for size exclusion chromatography in a buffer containing 20 mM HEPES pH 7.4, 400 mM NaCl, 5 mM MgCl_2_, 10% glycerol, and 1 mM DTT. Purified DDX3X was aliquoted, frozen in liquid nitrogen, and stored at -80°C (45).

### ZIKV 5’UTR RNA production

The 5’UTR of the ZIKV vRNA was recovered from plasmid pCC1-SP6-ZIKV (kindly provided by Dr. Andres Merits, University of Tartus), that encodes a complete viral genome (ZIKV isolate BeH819015, GenBank accession N° KX197192) by PCR. The PCR product was then used as a template for *in vitro* transcription using the mMessage mMachine SP6 kit following the manufacturer’s instructions (36). RNA was precipitated in the presence of 2.5 M LiCl and absolute ethanol, centrifuged at 16000g for 30 min at 4°C, washed with 70% ethanol, and resuspended in 50 µL of nuclease-free water. RNA integrity was monitored by electrophoresis on agarose gels. Finally, RNA was treated with Turbo DNase following manufacturer’s instructions. This RNA was used for EMSA and microscale thermophoresis. 5’UTR sequence: 5’*AGUUGUUGAUCUGUGUGAAUCAGACUGCGACAGUUCGAGUUUGAAGCGAAAGCUAGCAACAGUAUCAACAGGUUUUAUUUUGGAUUUGGAAACGAGAGUUUCUGGUCAUGAAAAACCCAAAAAAGAAAUCCGGAGGAUUCCGGAUUGUCAAUAUGCUAAAACGCGGAGUAGCCCGUGUGAGCCCCUUUGG* ‘3.

### Electrophoretic Mobility Shift Assay

Increasing concentrations of purified DDX3X (132–607) were used, which were mixed with a fixed amount of RNA (100 nM of 5’UTR region of the viral RNA) for 30 minutes at room temperature in RNA binding buffer (1 mM HEPES, 150 mM NaCl, 3 mM EDTA, 2.5 mM MgCl_2_, 1 mM DTT) (46). Then, 8 µL of the sample was taken and mixed in a 1:1 ratio with the loading buffer for EMSA (40% sucrose, 0.005% bromine phenol blue). From the mixture, 8 µL was loaded onto a TBE/Acrylamide gel, and an electrophoretic run was performed at constant 100 V for 1 hour in 0.5X TBE buffer. Then, the gel was stained with SyBrGold (Cat. S11494, Invitrogen) for 30 minutes at room temperature and stirred to be recorded using the Q9 Alliance UVITEC-Cambridge digital photodocumenter.

### Microscale thermophoresis

RNA fluorescent labeling was carried out through oxidation reactions. In this process, 10 μM of RNA was incubated with sodium acetate (100 mM) and sodium iodate (100 μM) for 90 minutes at room temperature. Subsequently, the oxidized RNA was precipitated using 100% ethanol and molecular grade glycogen (ThermoFischer Scientific). To perform conjugation reaction at the 3’ end of the RNA, fluorescein 5-thiosemicarbazide (FTSC) was used as a fluorescent molecule; the oxidized RNA was incubated with a labeling solution containing 1.5 mM of FTSC and 100 mM of sodium acetate solution. The labeled RNA (vRNA-FTSC) was precipitated, quantified and stored at −20 °C until use (47). For microscale thermophoresis (MST) experiments, serial dilutions of purified DDX3X (132–607) were incubated with a constant amount of vRNA-FTSC (100 nM) in a 1:1 ratio in MST buffer (20 mM HEPES pH 7.4, 150 mM NaCl, 2.5 mM MgCl_2_, 5% glycerol, 0.005% Tween). After incubation, samples were added to premium coated capillaries (NanoTemper Technologies) and subsequently subjected to MST analysis. Finally, the fluorescence values obtained were adjusted according to a unique binding model and the dissociation constant between 5’UTR of the and DDX3X (132–607) was determined using the MO.Affinity v2.3 software (NanoTemper Technologies)(48).

### Reporter ZIKV mRNA production and *in vitro* translation

Reporter ZIKV mRNA was produced from plasmid pCC1-SP6-ZIKV-NanoLuc (kindly provided by Dr. Andres Merits, University of Tartus), that encodes a complete viral genome that express a capsid protein fused to nanoluciferase enzyme (ZIKV isolate BeH819015, GenBank accession n° KX197192) by PCR. The PCR product was then used as a template for *in vitro* transcription using the mMessage mMachine SP6 kit following the manufacturer’s instructions (36). RNA was precipitated in the presence of 2.5 M LiCl and absolute ethanol, centrifuged at 16000 g for 30 min at 4°C, washed with 70% ethanol, and resuspended in 50 µL of nuclease-free water. RNA integrity was monitored by electrophoresis on agarose gels. Then, reporter RNA was treated with Turbo DNase following manufacturer’s instructions. *In vitro* translation reactions were carried out in nuclease-treated Flexi rabbit reticulocyte lysate (RRL) (Cat. L4960, Promega Corporation) at 35% vol/vol, supplemented with 10 μM amino acids (Promega Corporation) and 5 U/μL Ribolock RNase inhibitor (ThermoFisher Scientific). Reporter Zika mRNA was used at a final concentration of 0.5 pmol with different molar concentrations of DDX3X protein (0:1 to 10:1 DDX3X/RNA molar ratio). Translation reactions were incubated at 30°C for 90 min and luciferase activity was measured using the Nano-Glo® Luciferase Assay System (Promega Corporation) on Sirius Single Tube Luminometer (Berthold Detection Systems, GmbH, Pforzheim, Germany). Data are expressed as a percentage of the Relative Luciferase Activity Units (RLU) of control condition.

### Computational modeling

The computational modeling of the interaction between DDX3X and ZIKV RNA 5’UTR structure was performed using a combination of docking and classical molecular dynamics methods. First, we prepared the DDX3X receptor using the crystallographic structure at the pre-unwound state deposited in the Protein Data Bank (PDB ID: 6OF5)(49, 50). The CHARMM-GUI server (51) was used to apply corrections for discontinuities, add hydrogens and assign protonation states (pH 7.2) to the protein. Then, we used the prepared structure of the DDX3X and the 5’UTR sequence as inputs for protein-RNA docking through the HDOCK server (52). The three-dimensional structure of the RNA was generated during the same docking procedure conducted by HDOCK. A total of 100 protein-RNA conformations were generated. The structure with the best score provided by the software was selected, as it exhibited an RNA structure at the active site like DDX3X-dsRNA crystal complex (6O5F). Classical molecular simulation of each system was performed using NAMD 2.14 software (53). A TIP3P solvation box was defined at 15 Å from the complex. A concentration of 0.15 M of NaCl ions was added to ensure the net neutral charge of the system, which is for the Particle Mesh Ewald method (54) and to mimic physiological conditions. The 5’UTR and protein parameters were extracted from the CHARMM36m force field(55, 56). The simulation protocol for each solvated/neutralized system included: a) an initial minimization step comprising 20000 steps of the conjugate gradient algorithm, and b) 100 ns of NPT production simulation using the Particle-Mesh Ewald method (54) to calculate electrostatic contributions, and Langevin thermostat/Langevin barostat to maintain the temperature and pressure at 300 K and 101.325 kPa (1.0 atm), respectively. The equations of motion were integrated using a time step of 4 fs (57). All structural analyses were conducted using VMD software (58), ProLIF package (59) and NUCPLOT v.1.1.4 (60). The interaction between DDX3X and NS5 proteins was evaluated through molecular docking and molecular dynamics simulations. The full-length Zika virus NS5 protein structure was retrieved from the Protein Data Bank (PDB ID: 5M2X) (61) and prepared using the CHARMM-GUI server at pH 7.2. Subsequently, the prepared structures of DDX3X and NS5 were utilized as inputs for protein-protein docking evaluation through the HDOCK server. This process generated 100 protein-protein conformations. From these, the structure with the highest score was selected, as determined by the software, to serve as the initial structure for subsequent molecular dynamics analysis.

### Statistical analyses

Data are presented as mean ± SEM. The statistical analysis was performed using T-test or ANOVA (Dunnett post-test) in GraphPad Prism 9.0. *p*-value < 0.05 was considered statistically significant. All *p*-values are indicated in plots.

## RESULTS

### Induction of DDX3X expression during ZIKV replication in human microglia

ZIKV can efficiently infect and replicate in a wide range of cell types from different tissues including microglia, the major immune cells from the brain (33, 35). To investigate whether DDX3X could be a host factor involved in ZIKV replication in microglia, we analyzed the levels of the host RNA helicase in the immortalized adult human microglia cell line C20 infected for 24 hours with a Brazilian isolate of ZIKV (36). For comparisons, we also performed parallel infections of A549 (human lung epithelial) and HEK293T (human embryonic kidney) cells, two widely used cellular models for the study of flavivirus replication. Quantifications of intracellular ZIKV RNA by RT-qPCR suggest higher replication rates in A549 cells, followed by microglia and HEK293T cells (Fig. 1a). Analysis of the viral protease/RNA helicase NS3 by western blot revealed a similar trend, further indicating that A549 cells are the most susceptible and/or permissive cell line for this ZIKV isolate followed by microglia and HEK293T cells (Fig. 1b). Interestingly, we observed that DDX3X levels increased in human microglia but not A549 or HEK293T cells suggesting a cell-type specific role for this host RNA helicase in viral replication (Fig. 1b). Consistent with this idea, silencing of DDX3X resulted in a strong decrease in viral titers in microglia but not in A549 or HE293T cells indicating that DDX3X acts as a microglia-specific host factor for this ZIKV isolate (Fig. 1c).

**Figure 1.**
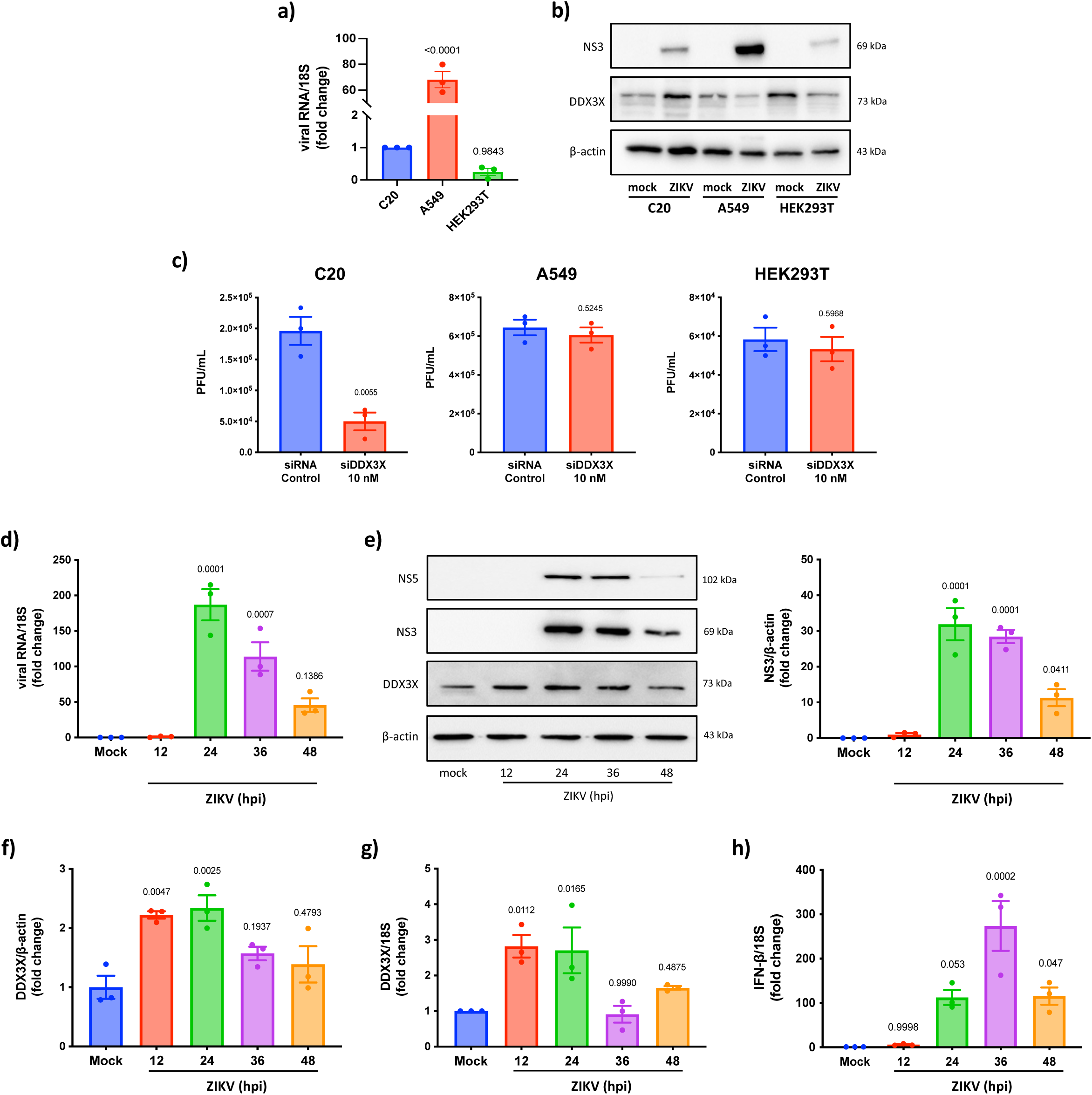
Induction of DDX3X expression during ZIKV replication in human microglia. (a) Intracellular total viral RNA detection by RT-qPCR at 24 hpi at MOI 3 in C20, A549 and HEK293T. 18S ribosomal RNA was used as housekeeping gene. Graph was truncated in Y axis. (b) Western blot of ZIKV infection in C20, A549 and HEK293T cells. Viral protease NS3, DDX3X and 𝛽-actin were evaluated at 24 hpi at MOI 3. (c) Viral titers in supernatant of DDX3X knockdown in C20 (left), A549 (middle) and HEK293T (right) cells determined by plaque assay. (d) Intracellular total viral RNA detection by RT-qPCR at 12, 24, 36 and 48 hpi at MOI 3 in human microglia cells. 18S ribosomal RNA was used as housekeeping gene. (e) Western blot of infection kinetics in C20 cells. Viral protease NS3, viral polymerase NS5, DDX3X and 𝛽-actin were evaluated at 12, 24, 36 and 48 hpi at MOI 3. Right panel, densitometric quantification of NS3 signal. (f) Densitometric quantification of DDX3X protein signal during ZIKV-infection kinetics in human microglia. (g) DDX3X mRNA detection by RT-qPCR at 12, 24, 36 and 48 hpi at MOI 3. 18S ribosomal RNA was used as housekeeping gene. (h) IFN-β mRNA detection by RT-qPCR at 24 hpi at MOI 3 in the three cells lines. 18S ribosomal RNA was used as housekeeping gene. Data are mean ± SEM from three independent experiments. Statistical significance was determined by One-way ANOVA and *p*-values are indicated in the plot.

To further characterize the induction of DDX3X by ZIKV in human microglia cells, we analyzed the levels of viral RNA, NS3, NS5 and DDX3X together with viral titers at 0, 12, 24, 36 and 48 hpi. Our results indicate that viral RNA reached a peak at 24 hpi that then decreased at 36 and 48 hpi (Fig. 1d). A similar trend was observed with NS3 and the RNA-dependent RNA polymerase NS5 (Fig. 1e), indicating that viral RNA and protein synthesis mostly take place during the first 24 hours of infection. Of note, viral titers increased over time suggesting sustained viral particle production and release during the time course analyzed (Supplementary Fig. 1a). Interestingly, we observed that the increase in DDX3X levels started at 12 hpi reaching a peak at 24 hpi to then return to basal levels at 48 hpi (Fig. 1f). DDX3X levels remained unchanged over time in uninfected cells indicating an increase due to viral replication (Supplementary Fig. 1b). RT-qPCR analyses of the DDX3X mRNA also showed and increase at 12-24 hpi that then reached basal levels suggesting that the augmented levels of DDX3X are due to an increase in its coding transcript (Fig. 1g). Such an induction of the DDX3X transcript occurred prior the induction of the IFN-β mRNA or the interferon stimulated gene retinoic acid inducible gene I (RIG-I), which peaked at 36 hpi (Fig. 1h and Supplementary Fig. 1c), indicating that induction of DDX3X during ZIKV infection might not be related to the type-I interferon response.

Together, these results indicate that ZIKV induces the expression of DDX3X at the levels of mRNA and protein during infection in human microglia to ensure efficient viral production.

### DDX3X is recruited to the viral replication compartment to promote ZIKV protein synthesis

Since DDX3X is a ubiquitously expressed protein that shuttles between the nucleus and the cytoplasm as a component of different ribonucleoprotein complexes and ZIKV replication takes place at viral replication compartments (VRC) located at the endoplasmic reticulum (62), we then sought to investigate whether DDX3X was recruited to the VRC during infection. As such, immunofluorescence staining, and confocal microscopy three-dimensional reconstitutions revealed that a fraction of DDX3X is recruited to the VRC during infection where it colocalizes with NS3 suggesting that the cellular RNA helicase might promote viral replication by regulating processes occurring at the replication sites (Fig. 2a). Consistent with this idea, silencing of DDX3X resulted in a potent reduction of viral proteins NS5 and NS3 indicating that DDX3X is required for the proper synthesis and/or accumulation synthesis of viral proteins (Fig. 2b). Moreover, fluorescent in situ hybridization and confocal microscopy three-dimensional reconstitutions showed that DDX3X colocalizes with the ZIKV RNA at the VRC suggesting a role for the RNA helicase in the synthesis of the viral polyprotein (Fig. 2c). We also evaluated whether DDX3X was required for protein synthesis during infection of human microglia by the DENV-2 virus. However, we could not observe an induction of DDX3X in DENV-2-infected microglia nor an effect of this RNA helicase on viral protein synthesis (Fig. 2d and Fig. 2e).

**Figure 2.**
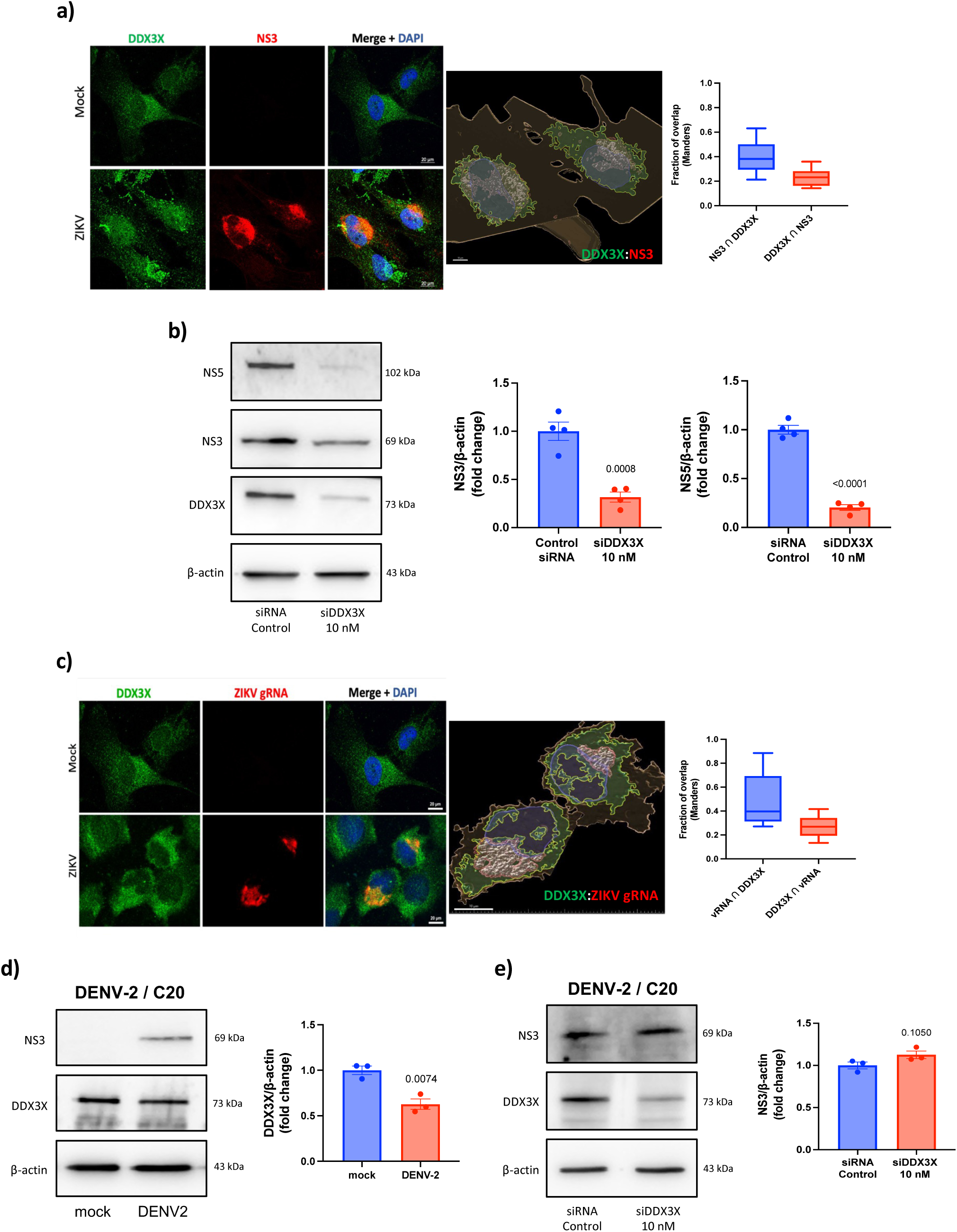
DDX3X is recruited to the viral replication compartment to promote ZIKV protein synthesis. (a) Confocal microscopy of infected C20 cells at 24 hpi at MOI 3. DDX3X in green, viral protease NS3 in red and nuclei was stained with DAPI. Inset correspond to the summatory of Z-stack reconstruction of colocalization volume using Imaris software. Right panel corresponds to Mander’s Overlap Coefficient (M) between NS3 and DDX3X signals. M1 of NS3 and DDX3X = 0.4 and M2 of DDX3X and NS3 = 0.247. (b) Western blot of DDX3X-knockdown C20 cells infected with ZIKV at 24 hpi. Viral polymerase NS5, viral protease NS3, DDX3X and 𝛽-actin were evaluated. Right panels correspond to densitometric quantification of NS3 and NS5 protein signals in DDX3X-knockdown conditions in C20 cells. (c) RNA fluorescent *in situ* hybridization coupled to indirect immunofluorescent microscopy (RNA FISH-IFI) was carried out in C20 cells infected for 24 hpi. Viral genome was detected using a digoxigenin-labeled RNA probe complementary to viral genome (in red), DDX3X in green and nuclei was stained with DAPI. Inset correspond to the summatory of Z-stack reconstruction of colocalization volume using Imaris software. The right panel correspond to Mander’s Overlap Coefficient (M) between vRNA and DDX3X signals. M1 of vRNA and DDX3X = 0.223 and M2 of DDX3X and vRNA = 0.264. (d) Western blot of DENV-2-infected C20 cells at 24 hpi. NS3, DDX3X and 𝛽-actin were evaluated. Right panel correspond to densitometric quantification of DDX3X protein signal in DENV-2-infected C20 cells. (e) Western blot of DDX3X-knockdown C20 cells infected with DENV-2 at 24 hpi. Viral protease NS3, DDX3X and 𝛽-actin were evaluated. Right panel correspond to densitometric quantification of NS3 protein signal in DDX3X-knockdown infection conditions. Data are mean ± SEM from three independent experiments. Statistical significance was determined by t-test, *p*-values are indicated in the plot.

Together, these data suggest that DDX3X is recruited to the ZIKV VRC to promote viral protein synthesis, and this effect was not observed in other flaviviruses, such as DENV-2.

### DDX3X binds the 5’UTR-Capsid region of the ZIKV vRNA to promote translation

Data from above showing that DDX3X is recruited to the VRC at the ER where it is required for the synthesis of the viral polyprotein together with the well-described roles of this RNA helicase in translation of selected mRNA targets carrying structured 5’ untranslated regions (5’UTR)(16, 39), strongly suggest that DDX3X could be involved in ZIKV vRNA translation. To demonstrate that DDX3X promotes translation by directly binding to the ZIKV vRNA, we performed EMSA using a constant concentration of an *in vitro* transcribed ZIKV-derived RNA containing the first 192 nucleotides of the viral genome that includes the entire 5’UTR and the first nucleotides of capsid coding region together with increasing amounts of purified recombinant DDX3X catalytic core (Supplementary Fig. 2a), that includes amino acids 132 to 607 (DDX3X^132-607^) (45). Through this assay we observed a marked change in the electrophoretic migration of the RNA at around to 2 µM of DDX3X (Fig. 3a). An EMSA using a globin-Renilla RNA previously shown to be independent of DDX3X activity(39) showed a change in the electrophoretic migration of the RNA between 5 and 7.5 µM of DDX3X, indicating that binding of DDX3X to the ZIKV RNA occurs with higher affinity (Supplementary Fig. 2b). We also performed microscale thermophoresis (MST) using a 3’-end FTSC labeled ZIKV-derived RNA (nt 1-192) to perform a similar binding experiment but with a higher sensitivity(47, 63). Following the same protocol used for the EMSA, a constant amount of ZIKV-derived RNA-FTSC was incubated with increasing concentrations of DDX3X. Processing of MST data suggests a dissociation constant (K_d_) of 11.7 µM for the DDX3X-ZIKV RNA complex (Supplementary Fig. 2c and Supplementary Fig. 2d), thus confirming the direct binding of DDX3X to the 5’UTR-Capsid region of the ZIKV vRNA as was described previously for Japanese encephalitis virus (48). Previous studies have proposed that the 5’UTR-capsid region of ZIKV and other flaviviruses folds into well-defined RNA motifs including stem-loop A (SLA), stem-loop B (SLB), capsid hairpin (cHP) and capsid coding region 1 (CCR1) loop, which are involved in the regulation of the translation-replication cycle (10, 64, 65). To gain further details on the 5’UTR and DDX3X interaction, we used the HDOCK server (52) together with previously reported SHAPE-Map data of the ZIKV vRNA^70^ to generate a three-dimensional model of the ZIKV 5’UTR-Capsid region. The obtained model preserved the highly conserved functional SL A, SLB, cHP and CCR1 domains (Fig. 3b). Then, we took advantage of the available structure of the DDX3X dimer(50) and conducted molecular simulations of the DDX3X-5’UTR pair with a production time of 100 ns under physiological conditions (Fig. 3c and Movie S1). This molecular simulation revealed that the most frequent interactions observed with monomer A were hydrogen bonds and salt bridges between Arg145 with A153 as well as Arg199 with A180 (Supplementary Fig. 2e). On the other hand, the primary interactions of DDX3X monomer B with ZIKV 5’UTR were observed between Lys451 and U188; Gly473 and G189 and Arg480 with G190 (Fig. 3c, inset and Supplementary Fig. 2e). Therefore, these molecular simulations showed that the 5’UTR exhibits a significant stability within the region spanning nucleotides 150-190 within CCR1 loop, primarily interacting with the D2-CTE region at the catalytic domain of monomer B within the DDX3X dimer (Figure 3c and Supplementary Figure 2e). Of note, this modeling agrees with the RNA duplex binding mechanism previously described by Song et al (50), where the DDX3X dimer establishes contacts with both strands of its RNA duplex target. Illustration of the DDX3X/5’UTR interactions by NUCPLOT showed that interactions were specific to nucleotides 150-166 (strand 1) in the CCR1 loop with monomer A and 172-189 (strand 2) with monomer B (Fig. 3d) and shows that DDX3X could stablish different kind of interactions with the viral RNA, including interactions with phosphate groups, backbone sugar as well as hydrogen bonds (Fig. 3d and Supplementary Fig. 2e). Additionally, Root-mean-square fluctuation (RMSF) analysis indicates an increased mobility in the DDX3X monomer B, associated with the initial accommodation of the system after molecular coupling (Supplementary Figure 2f and Movie S1 “DDX3X-5’UTR”). Conversely, the 5’UTR exhibits significant stability within the region spanning nucleotides 150-190, primarily interacting with the RNA helicase (Supplementary Figure 2f and Movie S1).

**Figure 3.**
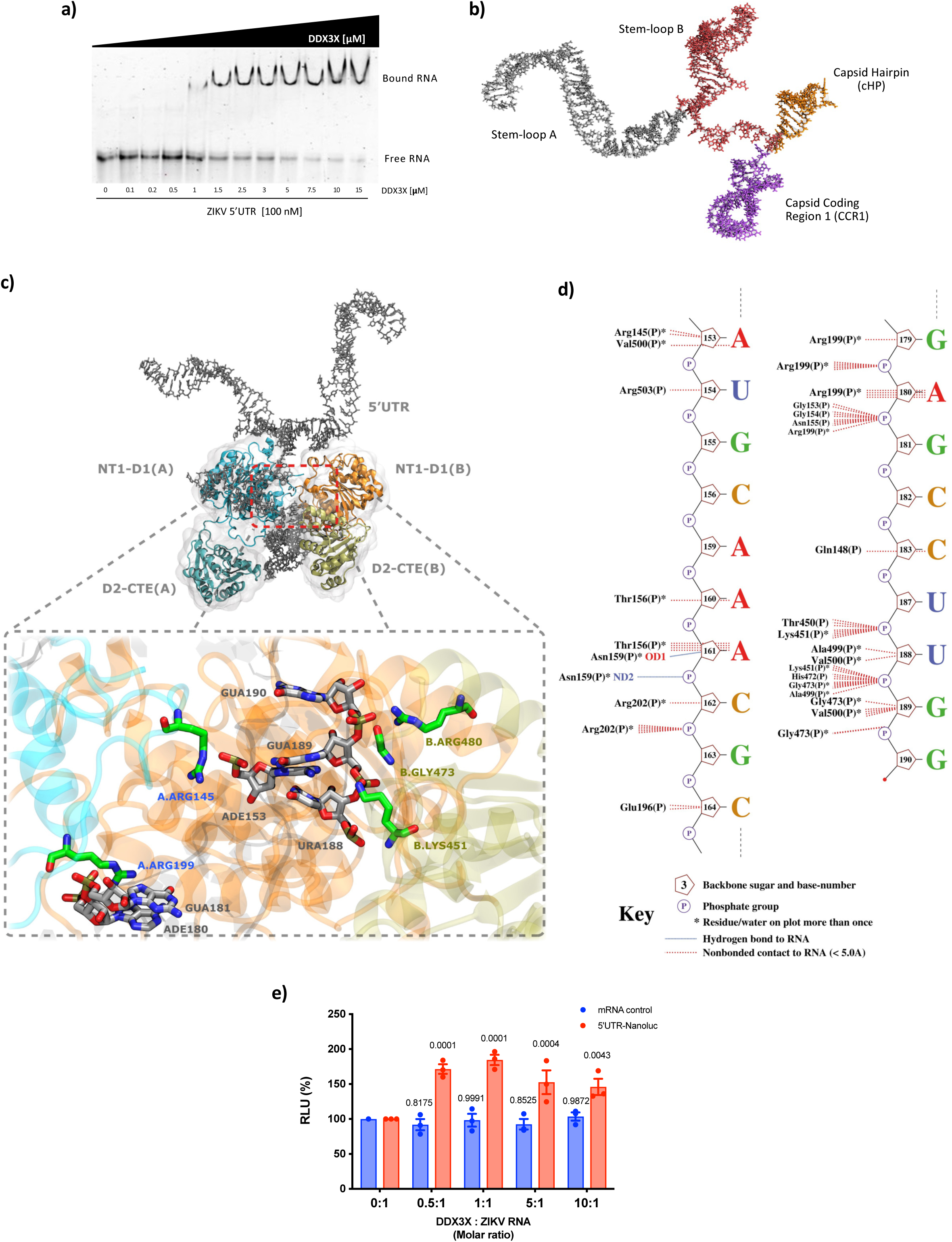
DDX3X binds the 5’UTR-Capsid region of the ZIKV vRNA to promote translation. (a) Electrophoretic mobility shift assay to evaluate DDX3X (132–607) and ZIKV 5’UTR interaction. Constant RNA concentration (100 nM) was incubated with increasing DDX3X (132–607) concentrations (0 to 15 μM). (b) Three-dimensional representation of ZIKV 5’UTR by simulation via HDOCK server. Conserved flaviviral functional regions are designated as follows: stem-loop A (gray), stem-loop B (red), capsid hairpin (cHP, yellow), and capsid coding region 1 (CCR1, purple). ZIKV 5’UTR sequence obtained from ZIKV isolate BeH819015 (GI: 975885966) was used. (c) Computational simulation of DDX3X-5’UTR interaction using HDOCK server. Representative structure of the last simulation frame with a close-up of the most relevant interactions during trajectory. Below, inset of DDX3X and 5’UTR principal interactions. Each DDX3X monomer was colored as follow: monomer A, with its N-terminal domain 1 (NT1-D1) in light blue and domain 2 at C-terminal end (D2-CTE) in blue. For monomer B, NT1-D1 in orange, and D2-CTE in yellow. (d) Summary of interactions between DDX3X and 5’UTR in the last frame of simulation using NUCPLOT software. (e) Cell-free translation in RRL was monitored using the viral reporter RNA and mRNA control incubated with increasing DDX3X:RNA molar ratio. Data are mean ± SEM from three independent experiments. Statistical significance was determined by One-way ANOVA compared against respective RNA reporter and t-test. *p*-values are indicated in the plot.

In this model, the accessibility to the cap structure is expected to be occluded by SLA while the initiation codon is embedded within SLB. The presence of these features characteristic of DDX3X targets within the 5’UTR-capsid region prompted us to investigate whether DDX3X was promoting translation of the ZIKV vRNA. For this, we generated a reporter ZIKV mRNA consisting of the 5’UTR-capsid coding region in-frame with the nanoluciferase (NanoLuc) reporter gene to program *in vitro* translation reactions in the rabbit reticulocyte lysate (RRL). In this cell-free system, the addition of DDX3X^132-607^ at an increasing molar ratio (protein: viral reporter RNA ratio 0:1 to 10:1) resulted in a significant increase in NanoLuc activity compared with a control reporter mRNA indicating that DDX3X can promote translation by acting on the ZIKV 5’UTR-Capsid region (Fig. 3e). Together, these data suggest that the DDX3X dimer binds to the 5’UTR-Capsid region probably at the CCR1 loop to promote viral RNA translation.

### DDX3X associates with the RdRp NS5 and interferes with viral RNA synthesis

Having demonstrated that DDX3X is required for viral protein synthesis together with evidence reporting the ability of the ZIKV RdRp NS5 to inhibit this process to displace the translation-replication cycle towards viral RNA replication (13), we sought to evaluate whether DDX3X was also involved in the regulation of viral RNA synthesis. Strikingly, we observed that DDX3X silencing resulted in a massive increase in vRNA levels (Fig. 4a). Furthermore, RNA FISH and microscopy analyses revealed that the vRNA accumulates, judged by an increase in MFI of ZIKV RNA, and remains at the VRC when DDX3X is silenced (Figure 4b), confirming vRNA accumulation observed by RT-qPCR. Such an effect was not observed when we analyzed the DENV-2 vRNA levels (Fig. 2e), which is consistent with the lack of impact of DDX3X silencing on DENV-2 NS3 accumulation.

**Figure 4.**
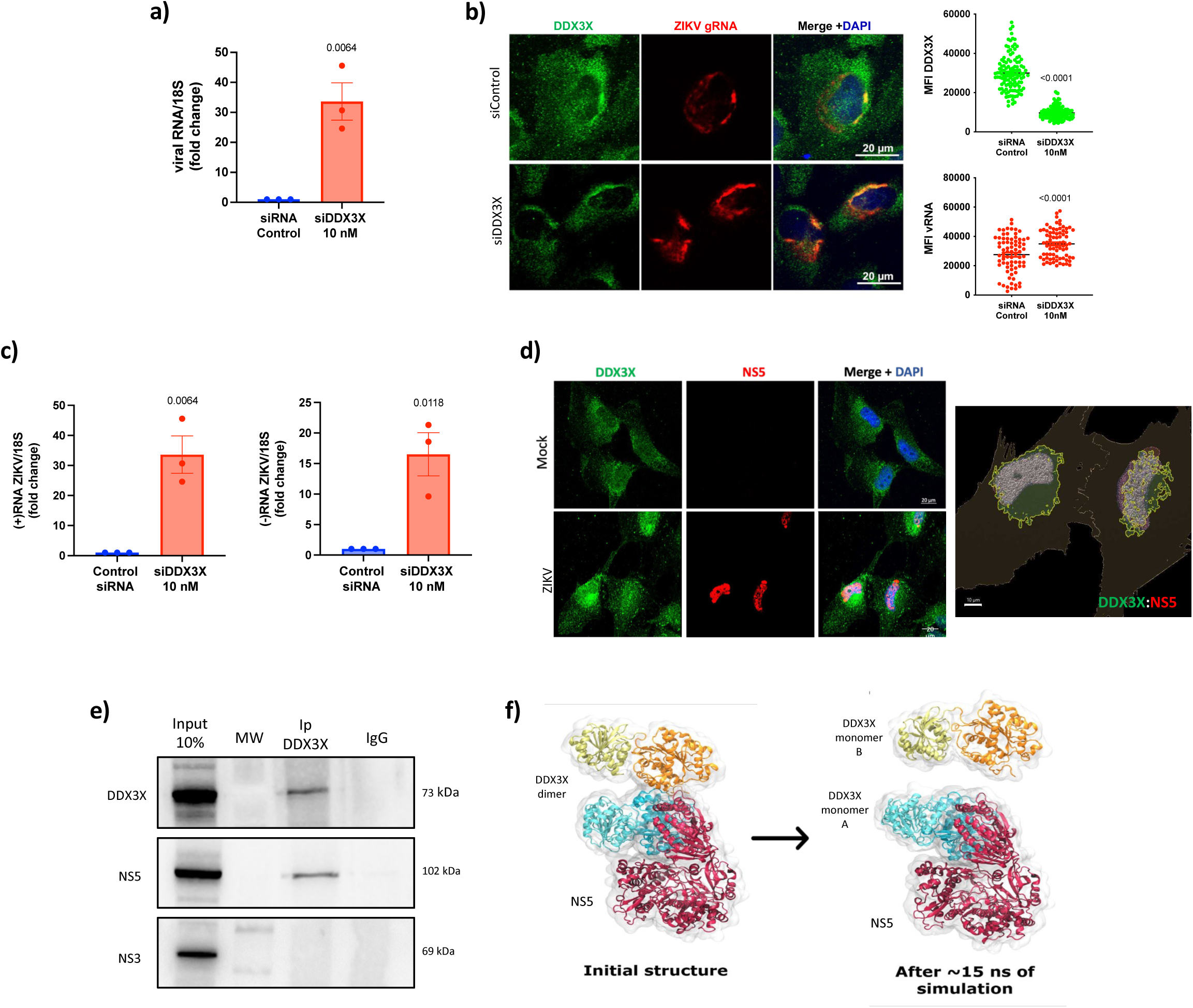
DDX3X associates with the RdRp NS5 and interferes with viral RNA synthesis. (a) Total viral RNA (vRNA) detection by RT-qPCR in DDX3X-knockdown conditions in C20 cells. 18S ribosomal RNA was used as housekeeping gene. (b) RNA FISH-IFI was carried out in DDX3X-knockdown C20 cells infected for 24 hpi. Viral genome was detected using a digoxigenin-labeled RNA probe (in red), DDX3X in green and nuclei was stained with DAPI. The right panels correspond to quantification of mean fluorescence intensity (MFI) of DDX3X and ZIKV gRNA signals in DDX3X knockdown microglia cells. Software ImageJ/Fiji was used. (c) Positive (left) and negative strand (right) viral RNA detection by strand specific RT-qPCR in DDX3X-knockdown conditions. 18S ribosomal RNA was used as housekeeping gene. (d) Confocal microscopy of infected C20 cells at 24 hpi. DDX3X in green, viral polymerase NS5 in red and nuclei was stained with DAPI. Inset correspond to the summatory of Z-stack reconstruction with Imaris software. (e) DDX3X co-immunoprecipitation was performed, viral polymerase NS5 and viral helicase NS3 was evaluated by western blot of immunoprecipitated fractions. (f) Simulation of the interaction between DDX3X-NS5. Disassembly of DDX3X dimer after 15 ns of DDX3X-NS5 system simulation. Stable interaction between DDX3X monomer A (blue) and NS5 (red) remains until the end of simulation. Statistical significance was determined by t-test compared against respective siRNA control group. *p*-values are indicated in the plot.

Since viral RNA replication by NS5 involves the synthesis of a negative strand intermediate from which the positive strand is produced, we conducted a strand-specific analysis of vRNA in control and DDX3X-silenced microglia. Interestingly, we observed that knocking down the host RNA helicase results in increased levels of both vRNA strands indicating that DDX3X might interfere with vRNA replication by NS5 (Fig. 4c). Immunostaining and confocal microscopy three-dimensional reconstitutions revealed that DDX3X and NS5 mostly colocalize in the nucleus (Fig. 4c), which is the main localization site described for this viral protein (66, 67). Moreover, immunoprecipitation analyses confirmed an interaction between DDX3X and NS5 but not NS3 (Fig. 4e). *In silico* simulations of the DDX3X-NS5 interaction shows a stable binding over the time between monomer A of DDX3X and NS5 (Fig. 4f). In detail, the interactions of these monomers maintain stable contacts mainly defined by hydrogen bonds and salt bridges between residues A.GLU216-NS5.LYS28, A.ASP354-NS5.ARG327, A.ASP354-NS5.LYS331, A.GLU388-NS5.327, A.GLU388-NS5.ARG741, A.ARG394-NS5.GLU112, and A.ASP398-NS5.ARG84 with frequencies over 80% (Fig. 4f and Supplementary Fig. 3a). Residues of NS5 interacting with DDX3X are located at the MTase domain, that includes SAM- and GTP-binding amino acids such as Lys28; Arg84 and Glu112. Additionally, DDX3X monomer A shows interactions with Arg327 and Lys331, belonging to thumb region of the RdRp domain. In the absence of an RNA sequence in the binding site, the DDX3X dimer exhibits a significant disassembly, where monomer B detaches from monomer A and monomer A remains attached to NS5, which might possibly interfere with NS5 functions in vRNA synthesis. This detachment and disassembly behavior is observed at an early stage of the simulation, around 15 ns (Fig. 4f, Supplementary Fig. 3b and Movie S2 “DDX3X-NS5”). This information allows us to understand structural effects of the DDX3X and NS5 association. Together, these results suggest that DDX3X associates with the ZIKV NS5 to prevent viral RNA synthesis.

### ATP-dependent and independent functions of DDX3X during ZIKV replication in human microglia

Since DDX3X is an ATP-dependent RNA helicase that also accomplishes functions independently its ATP binding/hydrolysis activities (20, 68), we finally sought to determine whether the functions of the cellular RNA helicase on vRNA translation and replication described above required ATP. For this, we took advantage of RK-33, a small molecule shown to interfere with the ATP-hydrolysis activity of DDX3X (40), to analyze its effects on NS3/NS5 and vRNA levels, the DDX3X-vRNA association and viral titers. Interestingly, we observed that treatment of RK-33 resulted in a decrease in NS3 and NS5 levels but has a marginal impact on vRNA levels suggesting that the function of DDX3X in promoting vRNA translation occurs in an ATP-dependent manner (Fig. 5a and 5b). Since RNA binding and unwinding by DDX3X requires ATP binding/hydrolysis (50), we then analyzed the DDX3X-vRNA interaction through RNA immunoprecipitation from ZIKV-infected microglia treated or not with RK-33. Interestingly, we observed that DDX3X associates with vRNA and this association was impeded by treatment of RK-33 (Fig. 5c). Overall, the inhibition of vRNA binding and viral protein by RK-33 resulted in a potent inhibition of viral titers (Fig. 5d).

**Figure 5.**
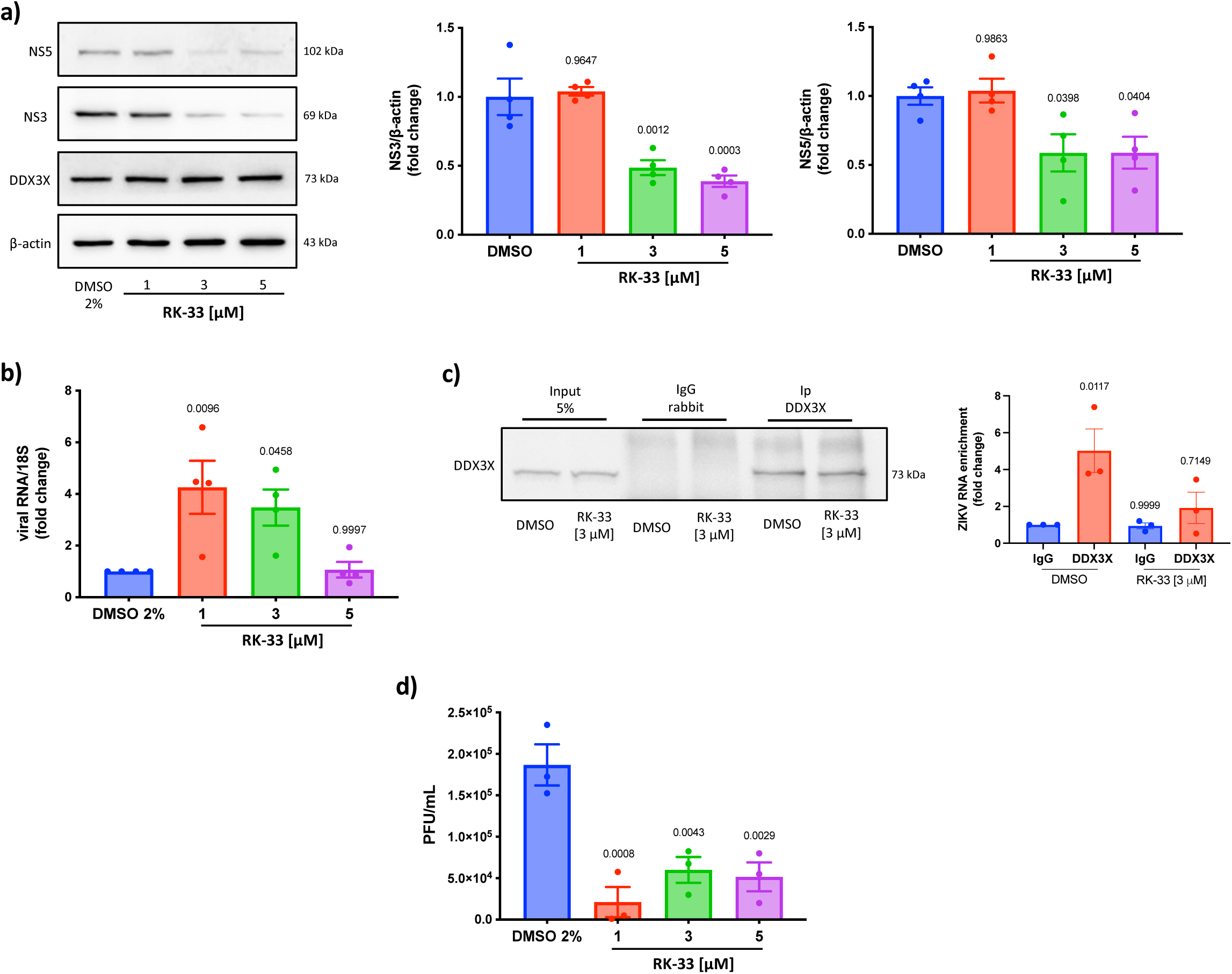
ATP-dependent and independent functions of DDX3X during ZIKV replication in human microglia. (a) Western blot of ZIKV-infected C20 cells treated with increasing RK-33 concentrations. NS5, NS3, DDX3X and 𝛽-actin were evaluated. The right panels correspond to densitometric quantification of NS3 and NS5 protein signals in RK-33 treatment conditions in C20 infected cells. (b) Total viral RNA (vRNA) detection by RT-qPCR in RK-33 treatment conditions. (c) CLIP-RTqPCR of DDX3X during ZIKV infection in microglia cells treated with DMSO or RK-33 at 3 μM for 24 hours. IgG fraction was used a specificity control of immunoprecipitation and input of DMSO treated cells was used for normalization and comparison. Data are mean ± SEM from three independent experiments. (d) Viral titers in supernatant of C20 infected cells treated with RK-33. Data are mean ± SEM from three independent experiments. Statistical significance was determined by One-way ANOVA and *p*-values are indicated in the plot.

Together, these results indicate that the functions of DDX3X in promoting vRNA translation requires ATP-binding/hydrolysis while its interference in vRNA synthesis occurs in an ATP-dependent manner.

## DISCUSSION

DDX3X is a cellular ATP-dependent RNA helicase that has a wide range of cellular functions, which includes cell cycle regulation, stress response, antiviral immune response, and mRNA metabolism (14, 15, 20). In mRNA metabolism, DDX3X participates in nuclear export, splicing, and translation initiation of a subset of mRNAs (15). Specifically, DDX3X is required for the translation of mRNAs that contain a highly structured 5’UTR close to the cap structure where its helicase activity is essential for the unwinding of RNA structures allowing 40S ribosomal subunit recruitment (16, 39, 69). Due to its multifaceted functions, several viruses including important human pathogens such as HCV, JEV, WNV and HIV-1, have been described to usurp the functions of DDX3X. As such, pharmacological inhibition of DDX3X has been shown to interfere with the replication of a wide range of RNA viruses in different cellular models and therefore, DDX3X has been proposed as a valuable target for the development of pan-antiviral agents against these viruses and other viruses with an impact on public health (70–72). Here, we provide evidence for a novel mechanism in which DDX3X acts as a dependency host factor for Zika virus replication in a human microglia cell line but not in human lung epithelial (A549) or embryonic kidney (HEK293T) cells. In agreement with this notion, we detected an increase in the DDX3X mRNA and protein levels during ZIKV infection in human microglia but not in A549 or HEK293T cells. Such an increase of DDX3X was readily detected at 12 hpi prior the peak of viral protein and RNA synthesis, which occurred at 24 hpi. Although DDX3X has been proposed as an interferon-stimulated gene (73), our data suggest that, at least in our model, an increase in the levels of DDX3X occurs independently from the type-I interferon response triggered by infection. In this regard, it is tempting to speculate that DDX3X levels increase early during infection to be recruited at the VRC in the ER and ensure efficient protein synthesis prior the induction of the antiviral interferon response, which has a peak at 36 hpi as evidenced by an increase in the INF-β mRNA and the protein levels of RIG-I.

Mechanistically, we provide evidence for a direct binding of DDX3X to the vRNA, most probably at the CCR1 stem-loop, to promote viral protein synthesis. Since the polyprotein initiation codon is embedded into the CCR1 stem-loop, it is tempting to speculate that DDX3X may facilitate the accessibility of the AUG codon to the 43S preinitiation complex through the unwinding of secondary structures present at the CCR1. This idea is further supported by the requirement of the ATP-binding/hydrolysis activity of DDX3X to ensure ZIKV protein synthesis. Whether DDX3X promotes cap-dependent or IRES-driven ribosome recruitment onto the ZIKV RNA requires further investigation.

An intriguing observation from our study was the massive accumulation of the positive and negative strands of the vRNA in the absence of DDX3X, which suggested a positive role for the host RNA helicase in vRNA translation but a negative regulation of vRNA synthesis by the RdRp NS5. Interestingly, NS5 was described to bind stem-loop A at the 5’UTR to promote the vRNA circularization required for the inhibition of translation and the promotion of RNA replication (12, 13). This apparent role of DDX3X in counteracting the activity of NS5 prompted us to investigate a possible interaction between both proteins. As such, we confirmed the interaction between DDX3X and NS5 during infection, and in silico molecular simulations suggest that DDX3X would interfere with the RdRp activity of NS5 by a direct binding to key residues in the thumb domain. In agreement with this idea, we observed that inhibiting the ATP-binding/hydrolysis activity of DDX3X has no effects on vRNA synthesis as observed upon DDX3X silencing further suggesting an interference mediated by protein-protein interactions and not by a processive event involving ATP hydrolysis.

Collectively, our results provide evidence for specific interactions between ZIKV and human microglia, which maintain the homeostasis of the translation-replication cycle of the vRNA ensuring efficient production of viral particles and the spread the infection. Whether DDX3X is also required for efficient replication in other myeloid cells such as monocytes, a key target cell for the virus in blood (74, 75), requires further investigation.

## DECLARATIONS

### Ethics approval and consent to participate

Not applicable.

### Consent for publication

Not applicable.

### Availability of data and materials

### Competing interests

The authors declare no conflict of interest.

## Funding

Work at the Laboratory of Molecular and Cellular Virology is funded by ANID - Chile through the FONDECYT Program grants N° 1230102 (to R.S.-R.), N° 1251218 (to F.V.-E), N° 1231275 (to M.L.A.), N° 1210736 (to M.L.-L.) and N° 11230976 (to M.Z.-B). R.S.-R, F.V.-E., M.L.A. and M.L.-L. holds funding from the ANID-ICM, grant N° ICN2021_045. T.H.-D. got an ANID Doctoral Fellowship N° 2019-21191987.

## Authors’ contributions

T.H.-D., A.G.-P, S.G.-O., A.O.-A, A.G.-A performed experiments and data analysis. C.R.-F. and M.Z.-B. performed *in silico* analyses. D.L.-P, L.C.-P. and F.C.G. performed RNA-FISH, immunofluorescences and image analysis. B.R.-A. and M.L.-L. performed translation experiments. T.H.-D. and T.C. helped with *in vitro* binding assays. T.H.-D. and R.S.-R. participated in conceptualization and writing. T.C., F.V.-E. and R.S.-R. corrected the manuscript and provided funding. All authors have read and agreed to the published version of the manuscript.

## Acknowledgements

Authors wish to thank Dr. Andres Merits (University of Tartu, Estonia) for providing the Zika virus reverse genetic systems and the NS3 and NS5 antibodies. Authors also wish to thank Dina Silva and Marisol Tapia (Universidad de Chile) for her technical support.

---

**Supplementary Figure 1.**

(a) Viral titers released to supernatant during infection kinetics. C20 cells were infected for 12, 24, 36, 48 hours and supernatant were collected. Viral titers were determined by plaque assay. (b) Western blot of DDX3X and at 0, 24 and 48 hours after seed cells. Right panel corresponds to densitometric quantification of DDX3X signal in mock conditions at 24 and 48 hours. (c) Western blot of infection kinetics in C20 cells. RIG-I and 𝛽-actin were evaluated at 12, 24, 36 and 48 hpi at MOI 3. Right panel correspond to densitometric quantification of RIG-I protein signal during infection kinetics. Data are mean ± SEM from three independent experiments. Statistical significance was determined by One-way ANOVA and *p*-values are indicated in the plot.

**Supplementary Figure 2.**

(a) Purification of recombinant DDX3X^132–607^. SDS-PAGE indicating the size exclusion chromatography (SEC)-purified DDX3X^132–607^. Lines 25 to 30 (in red) were collected and concentrated at 75 μM and storage at -80°C until use. (b) Electrophoretic mobility shift assay to evaluate DDX3X^132-607^ and globin-Renilla RNA interaction as control. Constant RNA concentration (100 nM) was incubated with increasing DDX3X^132-607^ concentrations (0 to 15 μM). (c) Representative MST traces depicting the change in fluorescent migration of FTSC-5’UTR. (d) Binding curve of DDX3X^132-607^ and 5’UTR fluorescent labeled, adjusted one site binding fit model with his respective calculated K_d_. (e) Frequency and type of contacts between DDX3X monomers and 5’UTR categorized by chains and protein regions. NT1 correspond to N-terminal of DDX3X, CTE correspond to C-terminal end. Chain A correspond to monomer A in light blue and chain B is monomer B in orange-yellow. (f) Root-mean-square fluctuations (RMSF) profile per nucleotide of 5’UTR (upper panel) and per residue for DDX3X (lower panel) during simulation.

**Supplementary Figure 3.**

(a) Interaction frequencies and types of contacts between DDX3X dimer and NS5 through the simulation trajectory. (b) Root Mean Square Deviation of DDX3X separated by monomers and NS5.

**Supplementary Movie S1.**

Video of molecular dynamic simulation of interaction between DDX3X dimer and ZIKV 5’UTR. The simulation was performed using NAMD 2.14 software during 100 ns. Conditions of simulations were a TIP3P solvation box at 15 Å from the complex, 0.15 M of NaCl ions concentration, temperature of 300 K and 1 atm of pressure to mimic physiological conditions.

**Supplementary Movie S2.**

Video of molecular dynamic simulation of interaction between DDX3X dimer and NS5 protein. The simulation was performed using NAMD 2.14 software during 100 ns. Conditions of simulations were a TIP3P solvation box at 15 Å from the complex, 0.15 M of NaCl ions concentration, temperature of 300 K and 1 atm of pressure to mimic physiological conditions.

## LIST OF ABBREVIATIONS

CCR1: Capsid coding region 1
cHP: Capsid hairpin
CLIP: Cross-linking immunoprecipitation
CNS: Central nervous system
DENV: Dengue virus
DMEM: Dulbecco modified eagle medium
EMSA: electrophoretic mobility shift assay
ER: Endoplasmic reticulum
FISH: Fluorescent in situ hybridization
FTSC: Fluorescein-5-thiosemicarbazide
IFN-β: Interferon beta
ISG15: Interferon stimulated gene 15
MFI: Mean fluorescence 7intensity
MOI: Multiplicity of infection
MST: Microscale thermophoresis
NS: Nonstructural protein (1 to 5)
RIPA: Radioimmunoprecipitation buffer assay
RMSD: Root mean square deviation
RMSF: Root mean square fluctuation
RRL: Rabbit reticulocyte lysate
SEM: Standard error median
SL: Stem loop
UTR: Untranslated region
VRC: Viral replication compartment
vRNA: Viral RNA
WHO: World Health Organization
ZIKV: Zika virus

## REFERENCES

1. Kim CR, Counotte M, Bernstein K, Deal C, Mayaud P, Low N, Broutet N, Sexual Transmission of Zika virus Expert Meeting p. 2018. Investigating the sexual transmission of Zika virus. Lancet Glob Health 6:e24–e25.

2. Miner JJ, Diamond MS. 2017. Zika Virus Pathogenesis and Tissue Tropism. Cell Host Microbe 21:134–142.

3. Alonso-Palomares LA, Moreno-Garcia M, Lanz-Mendoza H, Salazar MI. 2018. Molecular Basis for Arbovirus Transmission by Aedes aegypti Mosquitoes. Intervirology 61:255–264.

4. Klase ZA, Khakhina S, Schneider Ade B, Callahan MV, Glasspool-Malone J, Malone R. 2016. Zika Fetal Neuropathogenesis: Etiology of a Viral Syndrome. PLoS Negl Trop Dis 10:e0004877.

5. Olagnier D, Muscolini M, Coyne CB, Diamond MS, Hiscott J. 2016. Mechanisms of Zika Virus Infection and Neuropathogenesis. DNA Cell Biol 35:367–72.

6. Sirohi D, Kuhn RJ. 2017. Zika Virus Structure, Maturation, and Receptors. J Infect Dis 216:S935–S944.

7. Sun G, Larsen CN, Baumgarth N, Klem EB, Scheuermann RH. 2017. Comprehensive Annotation of Mature Peptides and Genotypes for Zika Virus. PLoS One 12:e0170462.

8. Ming GL, Tang H, Song H. 2016. Advances in Zika Virus Research: Stem Cell Models, Challenges, and Opportunities. Cell Stem Cell 19:690–702.

9. Barrows NJ, Campos RK, Liao KC, Prasanth KR, Soto-Acosta R, Yeh SC, Schott-Lerner G, Pompon J, Sessions OM, Bradrick SS, Garcia-Blanco MA. 2018. Biochemistry and Molecular Biology of Flaviviruses. Chem Rev 118:4448–4482.

10. Goertz GP, Abbo SR, Fros JJ, Pijlman GP. 2018. Functional RNA during Zika virus infection. Virus Res 254:41–53.

11. Cortese M, Goellner S, Acosta EG, Neufeldt CJ, Oleksiuk O, Lampe M, Haselmann U, Funaya C, Schieber N, Ronchi P, Schorb M, Pruunsild P, Schwab Y, Chatel-Chaix L, Ruggieri A, Bartenschlager R. 2017. Ultrastructural Characterization of Zika Virus Replication Factories. Cell Rep 18:2113–2123.

12. Sanford TJ, Mears HV, Fajardo T, Locker N, Sweeney TR. 2019. Circularization of flavivirus genomic RNA inhibits de novo translation initiation. Nucleic Acids Res 47:9789–9802.

13. Fajardo T, Sanford TJ, Mears HV, Jasper A, Storrie S, Mansur DS, Sweeney TR. 2020. The flavivirus polymerase NS5 regulates translation of viral genomic RNA. Nucleic Acids Res 48:5081–5093.

14. Ryan CS, Schroder M. 2022. The human DEAD-box helicase DDX3X as a regulator of mRNA translation. Front Cell Dev Biol 10:1033684.

15. Soto-Rifo R, Ohlmann T. 2013. The role of the DEAD-box RNA helicase DDX3 in mRNA metabolism. Wiley Interdiscip Rev RNA 4:369–85.

16. Calviello L, Venkataramanan S, Rogowski KJ, Wyler E, Wilkins K, Tejura M, Thai B, Krol J, Filipowicz W, Landthaler M, Floor SN. 2021. DDX3 depletion represses translation of mRNAs with complex 5’ UTRs. Nucleic Acids Res 49:5336–5350.

17. Saikruang W, Ang Yan Ping L, Abe H, Kasumba DM, Kato H, Fujita T. 2022. The RNA helicase DDX3 promotes IFNB transcription via enhancing IRF-3/p300 holocomplex binding to the IFNB promoter. Sci Rep 12:3967.

18. Shih JW, Wang WT, Tsai TY, Kuo CY, Li HK, Wu Lee YH. 2012. Critical roles of RNA helicase DDX3 and its interactions with eIF4E/PABP1 in stress granule assembly and stress response. Biochem J 441:119–29.

19. Samir P, Kesavardhana S, Patmore DM, Gingras S, Malireddi RKS, Karki R, Guy CS, Briard B, Place DE, Bhattacharya A, Sharma BR, Nourse A, King SV, Pitre A, Burton AR, Pelletier S, Gilbertson RJ, Kanneganti TD. 2019. DDX3X acts as a live-or-die checkpoint in stressed cells by regulating NLRP3 inflammasome. Nature 573:590–594.

20. Hernandez-Diaz T, Valiente-Echeverria F, Soto-Rifo R. 2021. RNA Helicase DDX3: A Double-Edged Sword for Viral Replication and Immune Signaling. Microorganisms 9.

21. Valiente-Echeverria F, Hermoso MA, Soto-Rifo R. 2015. RNA helicase DDX3: at the crossroad of viral replication and antiviral immunity. Rev Med Virol 25:286–99.

22. de Bisschop G, Ameur M, Ulryck N, Benattia F, Ponchon L, Sargueil B, Chamond N. 2019. HIV-1 gRNA, a biological substrate, uncovers the potency of DDX3X biochemical activity. Biochimie 164:83–94.

23. Soto-Rifo R, Rubilar PS, Ohlmann T. 2013. The DEAD-box helicase DDX3 substitutes for the cap-binding protein eIF4E to promote compartmentalized translation initiation of the HIV-1 genomic RNA. Nucleic Acids Res 41:6286–99.

24. Vesuna F, Akhrymuk I, Smith A, Winnard PT, Jr., Lin SC, Panny L, Scharpf R, Kehn-Hall K, Raman V. 2022. RK-33, a small molecule inhibitor of host RNA helicase DDX3, suppresses multiple variants of SARS-CoV-2. Front Microbiol 13:959577.

25. Yedavalli VS, Neuveut C, Chi YH, Kleiman L, Jeang KT. 2004. Requirement of DDX3 DEAD box RNA helicase for HIV-1 Rev-RRE export function. Cell 119:381–92.

26. Ariumi Y, Kuroki M, Abe K, Dansako H, Ikeda M, Wakita T, Kato N. 2007. DDX3 DEAD-box RNA helicase is required for hepatitis C virus RNA replication. J Virol 81:13922–6.

27. Ciccosanti F, Di Rienzo M, Romagnoli A, Colavita F, Refolo G, Castilletti C, Agrati C, Brai A, Manetti F, Botta L, Capobianchi MR, Ippolito G, Piacentini M, Fimia GM. 2021. Proteomic analysis identifies the RNA helicase DDX3X as a host target against SARS-CoV-2 infection. Antiviral Res 190:105064.

28. Brai A, Martelli F, Riva V, Garbelli A, Fazi R, Zamperini C, Pollutri A, Falsitta L, Ronzini S, Maccari L, Maga G, Giannecchini S, Botta M. 2019. DDX3X Helicase Inhibitors as a New Strategy To Fight the West Nile Virus Infection. J Med Chem 62:2333–2347.

29. Chahar HS, Chen S, Manjunath N. 2013. P-body components LSM1, GW182, DDX3, DDX6 and XRN1 are recruited to WNV replication sites and positively regulate viral replication. Virology 436:1–7.

30. Xu P, Shan C, Dunn TJ, Xie X, Xia H, Gao J, Allende Labastida J, Zou J, Villarreal PP, Schlagal CR, Yu Y, Vargas G, Rossi SL, Vasilakis N, Shi PY, Weaver SC, Wu P. 2020. Role of microglia in the dissemination of Zika virus from mother to fetal brain. PLoS Negl Trop Dis 14:e0008413.

31. Christian KM, Song H, Ming GL. 2019. Pathophysiology and Mechanisms of Zika Virus Infection in the Nervous System. Annu Rev Neurosci 42:249–269.

32. Wang J, Liu J, Zhou R, Ding X, Zhang Q, Zhang C, Li L. 2018. Zika virus infected primary microglia impairs NPCs proliferation and differentiation. Biochem Biophys Res Commun 497:619–625.

33. Chen Z, Zhong D, Li G. 2019. The role of microglia in viral encephalitis: a review. J Neuroinflammation 16:76.

34. Li C, Wang Q, Jiang Y, Ye Q, Xu D, Gao F, Xu JW, Wang R, Zhu X, Shi L, Yu L, Zhang F, Guo W, Zhang L, Qin CF, Xu Z. 2018. Disruption of glial cell development by Zika virus contributes to severe microcephalic newborn mice. Cell Discov 4:43.

35. Martinez Viedma MDP, Pickett BE. 2018. Characterizing the Different Effects of Zika Virus Infection in Placenta and Microglia Cells. Viruses 10.

36. Mutso M, Saul S, Rausalu K, Susova O, Zusinaite E, Mahalingam S, Merits A. 2017. Reverse genetic system, genetically stable reporter viruses and packaged subgenomic replicon based on a Brazilian Zika virus isolate. J Gen Virol 98:2712–2724.

37. Medina F, Medina JF, Colon C, Vergne E, Santiago GA, Munoz-Jordan JL. 2012. Dengue virus: isolation, propagation, quantification, and storage. Curr Protoc Microbiol Chapter 15:Unit 15D 2.

38. Baer A, Kehn-Hall K. 2014. Viral concentration determination through plaque assays: using traditional and novel overlay systems. J Vis Exp doi:10.3791/52065:e52065.

39. Soto-Rifo R, Rubilar PS, Limousin T, de Breyne S, Decimo D, Ohlmann T. 2012. DEAD-box protein DDX3 associates with eIF4F to promote translation of selected mRNAs. EMBO J 31:3745–56.

40. Yang SNY, Atkinson SC, Audsley MD, Heaton SM, Jans DA, Borg NA. 2020. RK-33 Is a Broad-Spectrum Antiviral Agent That Targets DEAD-Box RNA Helicase DDX3X. Cells 9.

41. Frohlich A, Rojas-Araya B, Pereira-Montecinos C, Dellarossa A, Toro-Ascuy D, Prades-Perez Y, Garcia-de-Gracia F, Garces-Alday A, Rubilar PS, Valiente-Echeverria F, Ohlmann T, Soto-Rifo R. 2016. DEAD-box RNA helicase DDX3 connects CRM1-dependent nuclear export and translation of the HIV-1 unspliced mRNA through its N-terminal domain. Biochim Biophys Acta 1859:719–30.

42. Livak KJ, Schmittgen TD. 2001. Analysis of relative gene expression data using real-time quantitative PCR and the 2(-Delta Delta C(T)) Method. Methods 25:402–8.

43. Schindelin J, Arganda-Carreras I, Frise E, Kaynig V, Longair M, Pietzsch T, Preibisch S, Rueden C, Saalfeld S, Schmid B, Tinevez JY, White DJ, Hartenstein V, Eliceiri K, Tomancak P, Cardona A. 2012. Fiji: an open-source platform for biological-image analysis. Nat Methods 9:676–82.

44. Pereira-Montecinos C, Toro-Ascuy D, Ananias-Saez C, Gaete-Argel A, Rojas-Fuentes C, Riquelme-Barrios S, Rojas-Araya B, Garcia-de-Gracia F, Aguilera-Cortes P, Chnaiderman J, Acevedo ML, Valiente-Echeverria F, Soto-Rifo R. 2022. Epitranscriptomic regulation of HIV-1 full-length RNA packaging. Nucleic Acids Res 50:2302–2318.

45. Floor SN, Condon KJ, Sharma D, Jankowsky E, Doudna JA. 2016. Autoinhibitory Interdomain Interactions and Subfamily-specific Extensions Redefine the Catalytic Core of the Human DEAD-box Protein DDX3. J Biol Chem 291:2412–21.

46. Rio DC. 2014. Electrophoretic mobility shift assays for RNA-protein complexes. Cold Spring Harb Protoc 2014:435–40.

47. Zearfoss NR, Ryder SP. 2012. End-labeling oligonucleotides with chemical tags after synthesis. Methods Mol Biol 941:181–93.

48. Nelson C, Mrozowich T, Gemmill DL, Park SM, Patel TR. 2021. Human DDX3X Unwinds Japanese Encephalitis and Zika Viral 5’ Terminal Regions. Int J Mol Sci 22.

49. Berman HM, Westbrook J, Feng Z, Gilliland G, Bhat TN, Weissig H, Shindyalov IN, Bourne PE. 2000. The Protein Data Bank. Nucleic Acids Res 28:235–42.

50. Song H, Ji X. 2019. The mechanism of RNA duplex recognition and unwinding by DEAD-box helicase DDX3X. Nat Commun 10:3085.

51. Jo S, Kim T, Iyer VG, Im W. 2008. CHARMM-GUI: a web-based graphical user interface for CHARMM. J Comput Chem 29:1859–65.

52. Yan Y, Zhang D, Zhou P, Li B, Huang SY. 2017. HDOCK: a web server for protein-protein and protein-DNA/RNA docking based on a hybrid strategy. Nucleic Acids Res 45:W365–W373.

53. Phillips JC, Hardy DJ, Maia JDC, Stone JE, Ribeiro JV, Bernardi RC, Buch R, Fiorin G, Henin J, Jiang W, McGreevy R, Melo MCR, Radak BK, Skeel RD, Singharoy A, Wang Y, Roux B, Aksimentiev A, Luthey-Schulten Z, Kale LV, Schulten K, Chipot C, Tajkhorshid E. 2020. Scalable molecular dynamics on CPU and GPU architectures with NAMD. J Chem Phys 153:044130.

54. Sitlapersad RS, Thornton AR, den Otter WK. 2024. A simple efficient algorithm for molecular simulations of constant potential electrodes. J Chem Phys 160.

55. Denning EJ, Priyakumar UD, Nilsson L, Mackerell AD, Jr. 2011. Impact of 2’-hydroxyl sampling on the conformational properties of RNA: update of the CHARMM all-atom additive force field for RNA. J Comput Chem 32:1929–43.

56. Huang J, Rauscher S, Nawrocki G, Ran T, Feig M, de Groot BL, Grubmuller H, MacKerell AD, Jr. 2017. CHARMM36m: an improved force field for folded and intrinsically disordered proteins. Nat Methods 14:71–73.

57. Hopkins CW, Le Grand S, Walker RC, Roitberg AE. 2015. Long-Time-Step Molecular Dynamics through Hydrogen Mass Repartitioning. J Chem Theory Comput 11:1864–74.

58. Humphrey W, Dalke A, Schulten K. 1996. VMD: visual molecular dynamics. J Mol Graph 14:33-8, 27–8.

59. Bouysset C, Fiorucci S. 2021. ProLIF: a library to encode molecular interactions as fingerprints. J Cheminform 13:72.

60. Luscombe NM, Laskowski RA, Thornton JM. 1997. NUCPLOT: a program to generate schematic diagrams of protein-nucleic acid interactions. Nucleic Acids Res 25:4940–5.

61. Ferrero DS, Ruiz-Arroyo VM, Soler N, Uson I, Guarne A, Verdaguer N. 2019. Supramolecular arrangement of the full-length Zika virus NS5. PLoS Pathog 15:e1007656.

62. Mohd Ropidi MI, Khazali AS, Nor Rashid N, Yusof R. 2020. Endoplasmic reticulum: a focal point of Zika virus infection. J Biomed Sci 27:27.

63. Moon MH, Hilimire TA, Sanders AM, Schneekloth JS, Jr. 2018. Measuring RNA-Ligand Interactions with Microscale Thermophoresis. Biochemistry 57:4638–4643.

64. Fernandez-Sanles A, Rios-Marco P, Romero-Lopez C, Berzal-Herranz A. 2017. Functional Information Stored in the Conserved Structural RNA Domains of Flavivirus Genomes. Front Microbiol 8:546.

65. Liu ZY, Li XF, Jiang T, Deng YQ, Ye Q, Zhao H, Yu JY, Qin CF. 2016. Viral RNA switch mediates the dynamic control of flavivirus replicase recruitment by genome cyclization. Elife 5.

66. Ji W, Luo G. 2020. Zika virus NS5 nuclear accumulation is protective of protein degradation and is required for viral RNA replication. Virology 541:124–135.

67. Zhao Z, Tao M, Han W, Fan Z, Imran M, Cao S, Ye J. 2021. Nuclear localization of Zika virus NS5 contributes to suppression of type I interferon production and response. J Gen Virol 102.

68. De Colibus L, Stunnenberg M, Geijtenbeek TBH. 2022. DDX3X structural analysis: Implications in the pharmacology and innate immunity. Curr Res Immunol 3:100–109.

69. Wilkins KC, Schroeder T, Gu S, Revalde JL, Floor SN. 2023. Determinants of DDX3X sensitivity uncovered using a helicase activity in translation reporter. bioRxiv doi:10.1101/2023.09.14.557805.

70. Ma S, Mao Q, Weng S, Teng M, Luo J, Zhang K. 2025. DDX3X and virus interactions: functional diversity and antiviral strategies. Front Microbiol 16:1630068.

71. Brai A, Trivisani CI, Poggialini F, Pasqualini C, Vagaggini C, Dreassi E. 2022. DEAD-Box Helicase DDX3X as a Host Target against Emerging Viruses: New Insights for Medicinal Chemical Approaches. J Med Chem 65:10195–10216.

72. Winnard PT, Jr., Vesuna F, Raman V. 2021. Targeting host DEAD-box RNA helicase DDX3X for treating viral infections. Antiviral Res 185:104994.

73. Sharma N, Kessler P, Sen GC. 2023. Cell-type-specific need of Ddx3 and PACT for interferon induction by RNA viruses. J Virol 97:e0130423.

74. Foo SS, Chen W, Chan Y, Bowman JW, Chang LC, Choi Y, Yoo JS, Ge J, Cheng G, Bonnin A, Nielsen-Saines K, Brasil P, Jung JU. 2017. Asian Zika virus strains target CD14(+) blood monocytes and induce M2-skewed immunosuppression during pregnancy. Nat Microbiol 2:1558–1570.

75. Michlmayr D, Andrade P, Gonzalez K, Balmaseda A, Harris E. 2017. CD14(+)CD16(+) monocytes are the main target of Zika virus infection in peripheral blood mononuclear cells in a paediatric study in Nicaragua. Nat Microbiol 2:1462–1470.

